# Discoidan Domain Receptor 1 (DDR1) Tyrosine Kinase is Upregulated in PKD Kidneys But Does Not Play a Role in the Pathogenesis of Polycystic Kidney Disease

**DOI:** 10.1101/527614

**Authors:** Irfana Soomro, Aram Hong, Zhai Li, James S. Duncan, Edward Y. Skolnik

## Abstract

Tolvaptan is the only drug approved to slow cyst growth and preserve kidney function in patients with autosomal dominant polycystic kidney disease (ADPKD). However, its limited efficacy combined with significant side effects underscores the need to identify new and safe therapeutic drug targets to slow progression to end stage kidney disease. We identified DDR1 as receptor tyrosine kinase upregulated *in vivo* in 3 mouse models of ADPKD using a novel mass spectrometry approach to identify kinases upregulated in ADPKD. Previous studies demonstrating critical roles for DDR1 to cancer progression, its potential role in the pathogenesis of a variety of other kidney disease, along with the possibility that DDR1 could provide new insight into how extracellular matrix impacts cyst growth led us to study the role of DDR1 in ADPKD pathogenesis. However, genetic deletion of DDR1 using CRISPR/Cas9 failed to slow cyst growth or preserve kidney function in both a rapid and slow mouse model of ADPKD demonstrating that DDR1 does not play a role in PKD pathogenesis and is thus a not viable drug target. In spite of the negative results, our studies will be of interest to the nephrology community as it will prevent others from potentially conducting similar experiments on DDR1 and reinforces the potential of performing unbiased screens coupled with *in vivo* gene editing using CRISPR/Cas9 to rapidly identify and confirm new potential drug targets for ADPKD.

## Introduction

Dysregulation of kinases and the pathways they regulate play a prominent role in the pathogenesis of cyst growth in ADPKD^1, 2^. Moreover, pharmacologic inhibition of a number of different kinases up-regulated in ADPKD kidneys has been shown to slow cyst growth in animal models making kinase inhibitors among the most promising class of drugs for treating patients with ADPKD. We identified Discoidan Domain Receptor 1 (DDR1) as a previously unidentified kinase to be upregulated in 3 mouse models of ADPKD using a novel mass spectrometry approach^3-5^. DDR1 is a receptor tyrosine kinase (RTK) that mediates interaction with the extracellular matrix and is activated upon binding collagen^6, 7^. Recent evidence has indicated that DDR1 is up-regulated in various cancers and plays a role in tumor growth, progression, and invasion^8-11^. DDR1 has also been shown to play prominent roles in a number of kidney disease that include mouse models of Alports^12, 13^, obstructive uropathy^14^, the remnant kidney model of chronic kidney disease^15^ and nephrotoxic serum nephritis^16^ (**see the excellent review^17^).** Thus, based on these findings we entertained the possibility that DDR1 would play a prominent role in PKD pathogenesis and provide a link between the extracellular matrix and regulation of growth of cyst lining epithelia. However, despite the upregulation of both DDR1 protein and DDR1 kinase activity *in vivo* in mouse models of ADPKD, targeted deletion of DDR1 using CRISPR/Cas9 did not slow cyst growth or preserve kidney function in both an ““early rapid” and “late slow” mouse model of ADPKD.

## Materials and Methods

### Mass Spectrometry to identify kinases up and down-regulated in PKD kidneys

*Pax8rtTA*; *TetO-cre; PKD1*^*fl/fl*^ (PKD) or littermate *Pax8rtTA*; *PKD1*^*fl/fl*^ (control) mice were induced with 200 mg/kg of doxycycline in the diet X 2 weeks starting on PN day 28 and kidneys were harvested 5 weeks post induction. Kidneys were then lysed in lysis buffer containing kinase inhibitors and active kinases were affinity captured by passing lysates over multiplex inhibitor beads (MIB) containing a cocktail of phosphatase inhibitors as previously described^4, 5^. Bound kinases were then identified by LC separation followed by tandem mass spectrometry (LC-MS/MS)^4, 5^. Experiments were performed using 6 PKD and 6 WT kidneys isolated from independent animals (3 female and 3 male mice in each group).

### Genetic Deletion of DDR1 using CRISPR/Cas9

6 single-guide (sg) RNAs using Feng Zhang’s on line tool were screened in ES cells for their ability to create double strand breaks (DSB) in DDR1 and ultimately led to the identification of 2 sgRNAs complementary to regions in exons 2 and 3 of DDR1 that were most efficient in giving DSB and corresponded to with the following sequences: AGTAACGCAACCGATAGCTT and CTACCGCTGCCCGCCACAGC (see schematic Fig. 3a). RNA for these 2 sgRNAs were *in vitro* transcribed, purified, and microinjected together with Cas9 into *Pkd1*^*fl/fl*^ zygotes to generate *Ddr1*^*-/+*^*; Pkd1*^*f/fll*^ as previously described^18, 19^. Sequencing identified out-of-frame deletion of DDR1 in several mice. A similar strategy using CRISPR/Cas9 was used to generate DDR1^+/-^; *Pax8rtTA; TetO-cre; PKD1*^*fl/fl*^ mice.

### Mice

*Pkhd1-Cre;Pkd1*^*fl/fl*^ and *Pax8rtTA; TetO-cre; PKD1*^*fl/fl*^ mice have been previously reported^20^. *Aqp2-Cre* mice were purchased from Jackson Labs and crossed with *Pkd1*^*fl/fl*^ mice to generate *Aqp2-Cre; Pkd1*^*fl/+*^ mice as previously reported^21^. To generate *Pkhd1-Cre; DDR1*^*-/-*^*; Pkd1*^*fl/fl*^ mice, *DDR1*^*-/+*^*; Pkd1*^*fl/fl*^ mice generated using CRISPR/Cas9 were crossed with *Pkhd1-Cre; DDR1*^*-*^*/*^*+*^*;Pkd1*^*fl/+*^ (see schematic, Fig. 3a). PCR for the WT allele utilized primers 1, 2 (ATGCAGGACCGCACCATTCCTGA, GAGCTTCACACTTGGTGAGTACC), which yields a product of 256 bp. Deletion of parts of exons 2 and 3 and intervening intron amplifies a PCR product of 203 bp using primers 1, 3 (ATGCAGGACCGCACCATTCCTGA, CGCTCAGGCAATGACAGATGCTG). Offspring from this cross allowed us to assess mice with deletion of Pkd1 in DDR^-/-^ and DDR^+/-^ backgrounds. A separate cross of *Pkhd1-Cre;* DDR^+/+^*; Pkd1*^*fl/+*^ with DDR^+/+^*; Pkd1*^*fl/fl*^ was used to generate DDR^+/+^; *Pkhd1-Cre;Pkd1*^*fl/fl*^ mice. *Pkhd1-Cre* mice were sacrificed on day 22 at which time kidneys were harvested and analyzed as described below. Equal numbers of littermate female and male mice were studied. To assess whether dasatinib inhibited cyst growth, *Pkhd1-Cre;Pkd1*^*fl/fl*^ mice from the same litter were treated with vehicle or dasatinib by starting on day 12 and mice were sacrificed on day 22.

A similar strategy was used to generate *DDR1*^*-/-*^*; Pax8rtTA; TetO-cre; PKD1*^*fl/fl*^ and *DDR1*^*+/-*^*; Pax8rtTA; TetO-cre; PKD1*^*fl/fl*^ and *DDR1*^*+/+*^*; Pax8rtTA; TetO-cre; PKD1*^*fl/fl*^ mice. Cre was induced in these mice by adding 200 mg/kg of doxycycline in the diet X 2 weeks starting on PN day 28 and kidneys were harvested 14 weeks post induction as previously described^20^. All animals were used in accordance with scientific, humane, and ethical principles and in compliance with regulations approved by the New York University School of Medicine Institutional Animal Care and Use Committee.

### Cystic index and kidney immunohistochemistry

Kidneys were harvested, fixed in 4% paraformaldehyde for 4 hours at 4°C, and sagittal kidney sections were stained with hematoxylin and eosin. Sections were then photographed under the same magnification and cystic index was calculated using ImageJ analysis software on 2 sagittal sections/kidney as described^21, 22^. Cystic index was calculated as the cumulative cyst volume per total area of kidney^21, 22^.

Kidney immunohistochemistry was performed on ADPKD and normal human kidneys obtained from nephrectomy, and kidneys from *Pax8rtTA; Pkd1*^*fl/fl*^ (WT) and *Pax8rtTA; TetO-cre; PKD* ^*fl/fl*^ (PKD) mice following induction with doxycycline. Immunohistochemistry was performed on 4-µm formalin-fixed, paraffin-embedded kidney sections using antibodies as indicated. Chromogenic immunohistochemistry was performed on a Ventana Medical Systems Discovery XT platform with online deparaffinization, antigen retrieval and using Ventana’s reagents and detection kits. In brief, heat mediated antigen retrieval was performed using either CC1 (Tris-Borate-EDTA, pH 8.5) or RCC2 (Sodium Citrate pH6.0) as required. Endogenous peroxidase activity was blocked with 3% hydrogen peroxide. Primary antibodies were diluted in Dulbecco’s phosphate buffered saline (Life Technologies) 3 hours at 37°C and detected using anti-rabbit or anti-mouse HRP labeled multimers incubated for 8 minutes. The complex was visualized with 3,3 diaminobenzidene and enhanced with copper sulfate.

### Western Analysis

Kidneys harvested at time of sacrifice were flash frozen in liquid nitrogen, homogenized in lysis buffer, separated by SDS/PAGE, immunoblotted with primary antibody as indicated, and detected with a Li-cor IRdye secondary antibody as described^21^.

### Antibodies

The antibodies used included: Anti-DDR1, phospho-DDR1(Tyr792), MAPK, phospho-MAPK(Thr202/204), FAK, phospho-FAK(Tyr397), AKT, phosphor-AKT(ser473), Stat3, phospho-Stat3 (Tyr705) were purchased from Cell Signalling.

## Results

### Discoidan domain receptor 1 (DDR1) protein and kinase activity is upregulated in cyst lining epithelia *in vivo* in mouse and human ADPKD kidneys

To selectively enrich in an unbiased manner for active kinases in PKD kidneys, lysates from doxycycline-induced *Pax8rtTA*;*Pkd1*^*fl/fl*^ (control) and *Pax8rtTA; TetO-cre; Pkd1*^*fl/fl*^ mice (PKD kidneys) were passed over multiplex inhibitor beads and bound kinases were then identified by quantitative LC-MS (MIB) as previously described^4, 5^. One of the kinases identified was DDR1. We found that DDR1 protein and activity (assessed by anti-phospho-DDR1 antibodies) was increased in kidneys from 3 mouse models of ADPKD that included 2 “early, rapid” models in which Cre is driven by Pkhd1 (*Pkhd1-Cre; Pkd1*^*fl/fl*^) and aquaporin (*Aqp1-Cre; Pkd1*^*fl/fl*^, not shown) and in the “late, slow” *Pax8rtTA; TetO-Cre; Pkd1*^*fl/fl*^ model (**Fig. 1a**). In addition, DDR1 is expressed in mouse and human cyst lining epithelia (**Fig. 1b**).

**Figure 1.**
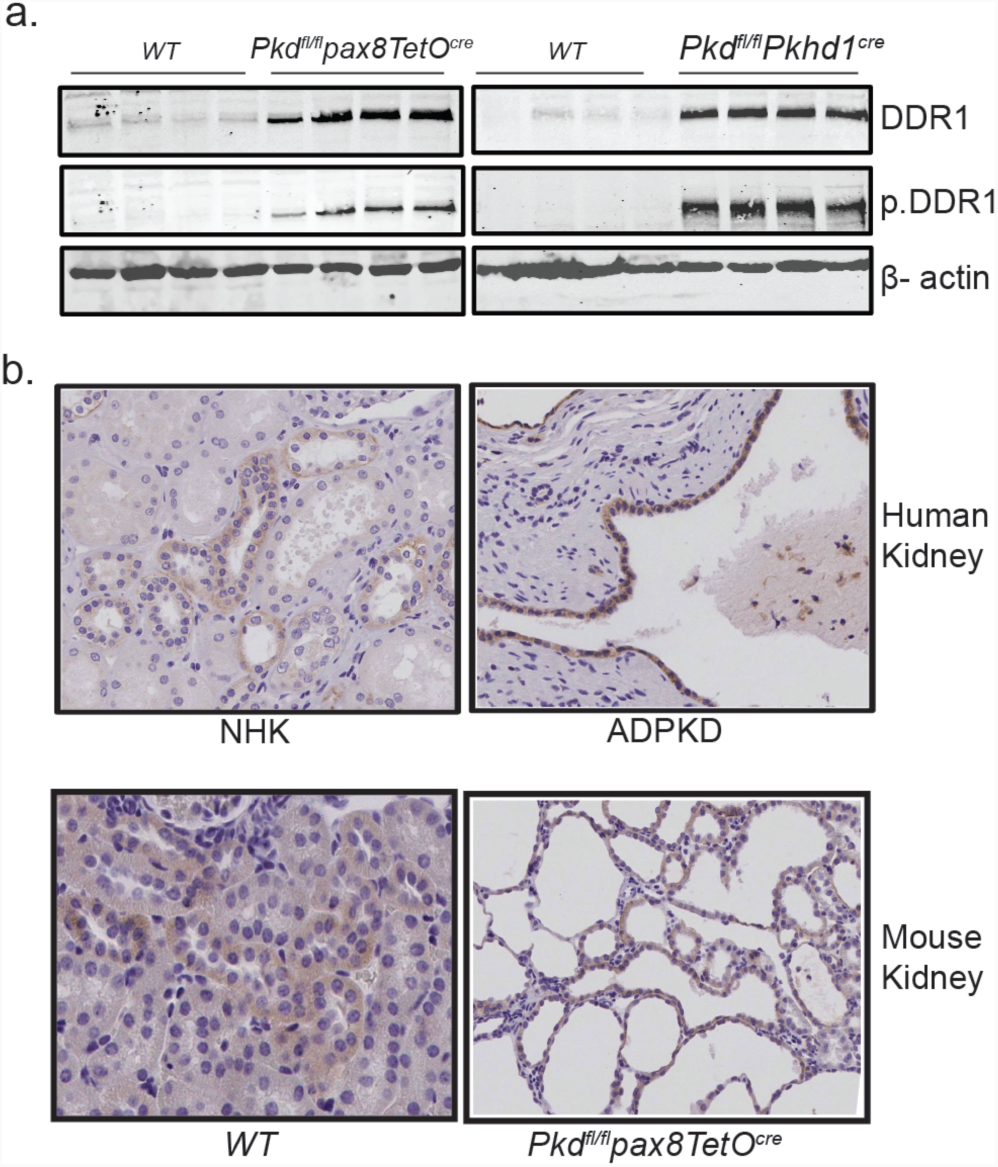
DDR1 protein and activity is increased in both early and late models of PKD. **(a)** Kidney tissue lysates from littermate and same sex *Pax8rtTA; TetO-cre; Pkd1fl/fl* and *Pax8rtTA; Pkd1fl/fl* (WT) treated with doxycycline and *Pkhd1-Cre;Pkd1fl/f* and *Pkd1fl/f* (WT) immunoblotted with anti-DDR1 (DDR1), anti-phospho-tyr-792-DDR1 antibodies (p.DDR1), and β-actin as a loading control. Results from 4 control and 4 PKD kidneys harvested from independent mice are shown. **(b)** Anti-DDR1immunohistochemistry: Upper panel, normal human (left) and ADPKD (right) kidneys; lower panel, *Pax8rtTA; Pkd1*^*fl/fl*^ (WT) (left) and *Pax8rtTA; TetO-cre; PKD* ^*fl/fl*^ (right) mice.

### Inhibiting DDR1 with a nonspecific DDR1 kinase inhibitor slowed cyst growth and preserved renal function in *Pkhd1-Cre; Pkd1*^*fl/fl*^ mice

Dasatinib is a small molecule nonspecfic inhibitor of DDR1. Treatment with dasatinib led to inhibition of DDR1 tyrosine phosphorylation, slowed cyst growth and preserved renal function in *Pkhd1-Cre; Pkd1*^*fl/fl*^ mice when compared with vehicle control treated mice, n=6 mice/group (**Fig. 2a-e**). To assess the signaling pathways affected by dasatinib treatment, kidneys from control and dasatinib treated mice were probed with various anti-phospho-antibodies. These studies demonstrated that dasatinib treatment not only inhibited activation of DDR1, but also Stat3 and AKT, while MAPK was not inhibited (**Fig. 2f**). Although these experiments provided a proof of concept that inhibiting DDR1 may slow cyst growth, dasatinib also inhibits other kinases^23-25^, such as Src family kinases and Kit.

**Figure 2.**
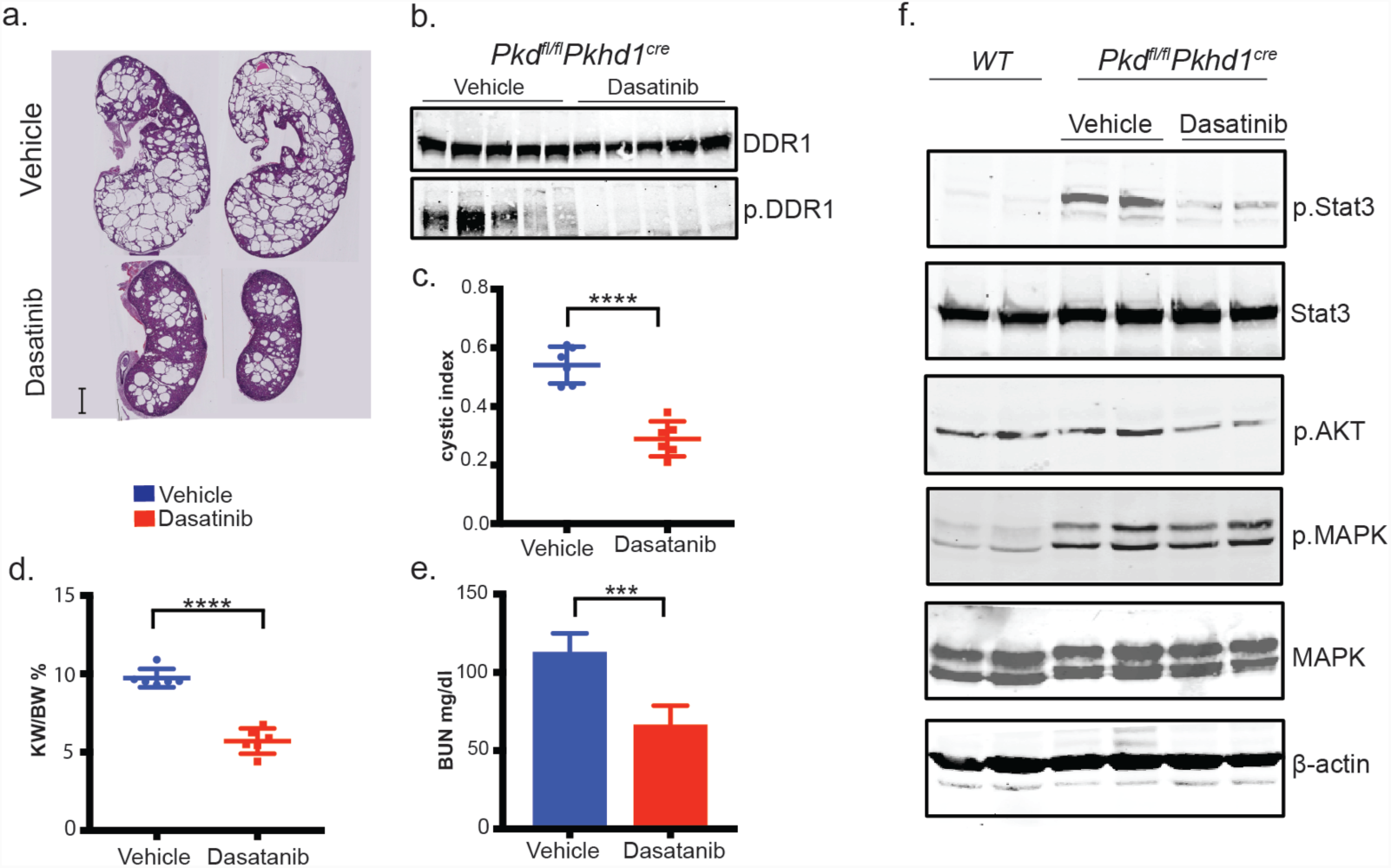
Pharmacologic inhibition of DDR1 with dasatinib slowed cyst growth in *Pkhd1-Cre;PKD1*^*fl/fl*^ mice and inhibits activation of STAT3 and AKT. Littermate same sex *Pkhd1-Cre;PKD1*^*fl/fl*^ mice were treated with vehicle control or dasatinib between days 12 and 22 when mice were sacrificed. **(a)** Representative histology from vehicle and dasatinib treated mice, **(b)** Kidney tissue lysate from vehicle (veh) and dasatinib treated mice immunoblotted with ddr1 antibody and p.Tyr 792 DDR1 antibody as shown **(add antibody to figure)**. (**c)** BUN **(d)** kidney/body weight ratio and **(e)** cystic index are shown. Cystic index was calculated using ImageJ analysis software on 2 sagittal sections/kidney as described^22, 26, 27^ and BUN ^28, 29^. **(f)** Lysates from kidneys from littermate same sex *Pkhd1-Cre;PKD1*^*fl/fl*^ mice treated with vehicle control or dasatinib were probed with antibodies as indicated. pStat3 –anti-phospho-Stat3, pAKT-anti-phospho (Thr 308) AKT), p.MAPK-anti-phospho-MAPK. Lysates were probed with β-actin as a loading control. Differences were evaluated by two-tailed *t*-tests (***=p<0.001, ****=p<0.0001).

### Genetic deletion of DDR1 fails to slow cyst growth and preserve renal function in *Pkhd1-Cre; Pkd1*^*fl/fl*^ mice

To definitively address whether DDR1 is critical for cyst growth, *DDR1*^*-/-*^ mice were generated by CRISPR/Cas9. DDR1 and PKD1 are both localized to chromosome 17 and therefore *Ddr1*^*-/+*^ mice cannot be crossed into *Pkd1*^*fl/fl*^ mice to generate *Ddr1*^*-/-*^; *Pkd1*^*fl/fl*^ mice. We used CRISPR/Cas9 to generate DDR1 knockouts in zygotes from *Pkd1*^*fl/fl*^ mice to generate a *Ddr1*^*-*^*; Pkd1*^*fl*^ chromosome 17 (**Fig. 3a**).

**Figure 3.**
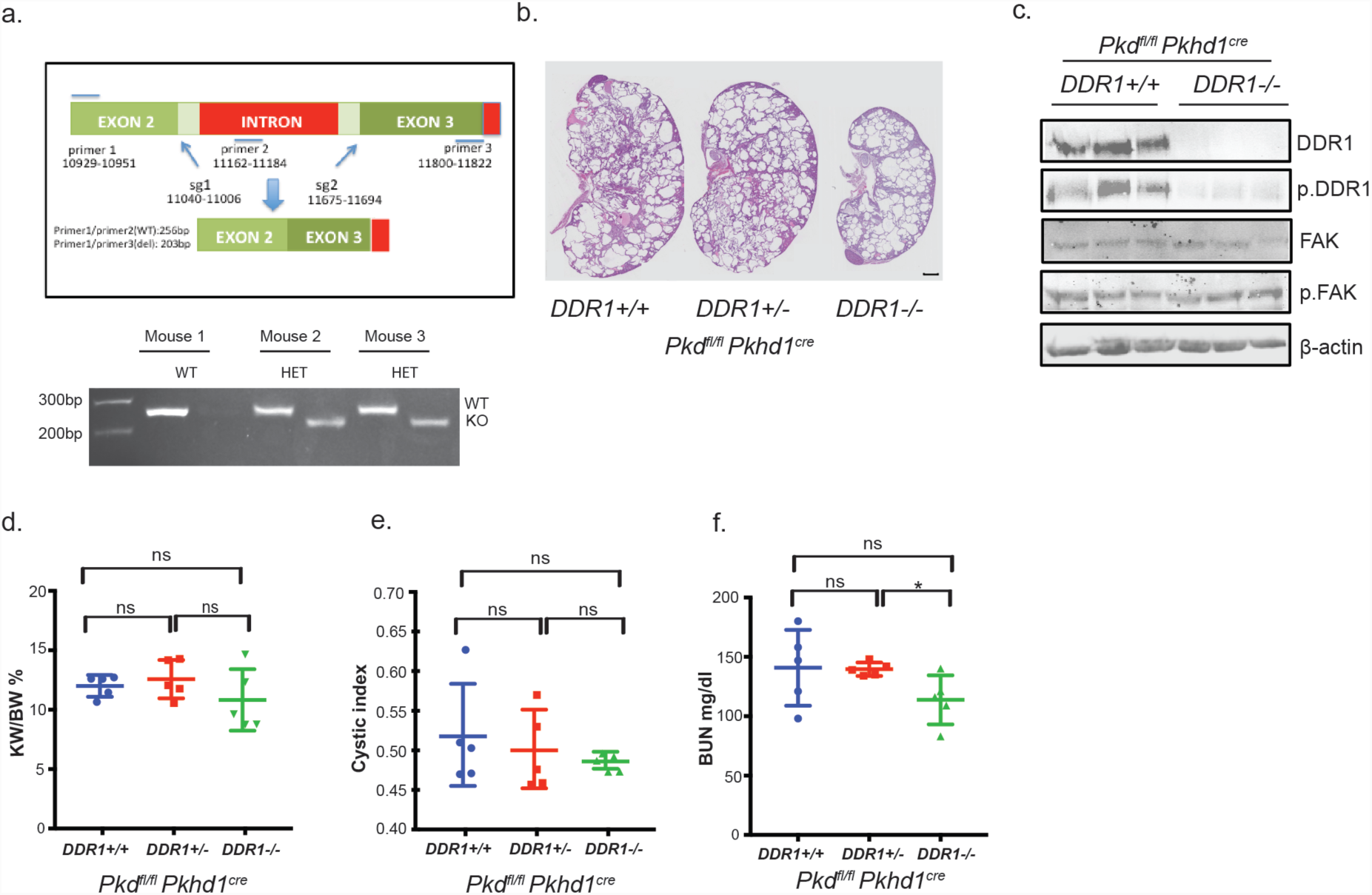
Generation of DDR1 KO mice and analysis of Pkhd1-Cre Pkd1 mice. (a) Schematic of DDR1 CRISPR/Cas9 KO. 2 sgRNAs to exon 2 and exon 3 were selected based on efficiency in ES cells. PCR for the WT allele utilized primers 1, 2, which yields a product of 256 bp. Deletion of parts of exons 2 and 3 and intervening intron amplifies a PCR product of 203 bp using primers 1, 3. Shown are PCR results from tail DNA isolated from DDR+/+ and DDR+/- mice using Primers 1,2 (256 bp, wt), and primers1,3 (203 bp, deleted). Under the conditions used primers 1,3 do not amplify the WT allele. PCR products were sequenced to demonstrate out of frame deletion. Numbering of bases labeled for sgRNAs and oligonucleotides corresponds to **gene ID:12305** and are shown in the Materials and Methods section. **(b)** Representative H&E staining of sagittal sections from DDR^+/+^; *Pkhd1-Cre;Pkd1*^*fl/fl*^, DDR^+/-^; *Pkhd1-Cre;Pkd1*^*fl/fl*^, and DDR^-/-^; *Pkhd1-Cre;Pkd1*^*fl/fl*^ mice. **(c)** Kidney lysates were probed as described in figure 2. Fak, focal adhesion kinase. pFak, p.FAK(Tyr397. **(d)** kidney/body weight ratio **(e)** cystic index and **(f)** (BUN are shown. Differences were evaluated by two-tailed *t*-tests (*=p<0.05, ns=not significant).

DDR1 and phospho-DDR1 (pDDR1) were absent from kidneys from *DDR1*^*-/-*^*; Pkhd1-Cre; Pkd1*^*fl/fl*^ mice confirming that our antibodies are specific for DDR1 and that we generated *DRR1*^*-/-*^ mice (**Fig. 3c**). However, we did not detect differences in kidney weight/body weight ratio (**Fig. 3b,d**), cystic index (**Fig. 3b,e**), or BUN (**Fig. 3f**) between *DDR1*^*-/-*^*; Pkhd1-Cre; Pkd1*^*fl/fl*^ and *DDR1*^*+/+*^*; Pkhd1-Cre; Pkd1*^*fl/fl*^ mice. These data indicate that the beneficial effect of dasatinib is via inhibition of a kinase(s) that is not DDR1.

### Genetic deletion of DDR1 also failed to slow cyst growth and preserve renal function in *Pax8rtTA; TetO-Cre; Pkd1*^*fl/fl*^ mice

In contrast to the *Pkhd1-Cre* model, deletion of PKD1 after postnatal day (PN) 14 leads to much slower cyst growth and loss of kidney function and, as a result, models in which PKD1 is inducibly deleted after PN day 14 are thought to more closely reflect disease in humans. We therefore also tested genetically if DDR1 played a role in cyst growth in the “slow late” *Pax8rtTA; TetO-Cre; Pkd1*^*fl/fl*^ model. These studies demonstrated that genetic deletion of DDR1 also did not slow cyst growth or preserve kidney function in *Pax8rtTA; TetO-Cre; Pkd1*^*fl/fl*^ mice; H&E staining of kidney sagittal sections, kidney/body weight ratio, and BUN were similar between *DDR1*^*-/-*^*; Pax8rtTA; TetO-Cre; Pkd1*^*fl/fl*^ and *DDR1*^*+/+*^*; Pax8rtTA; TetO-Cre; Pkd1*^*fl/fl*^ mice (**Fig. 4**).

**Figure 4.**
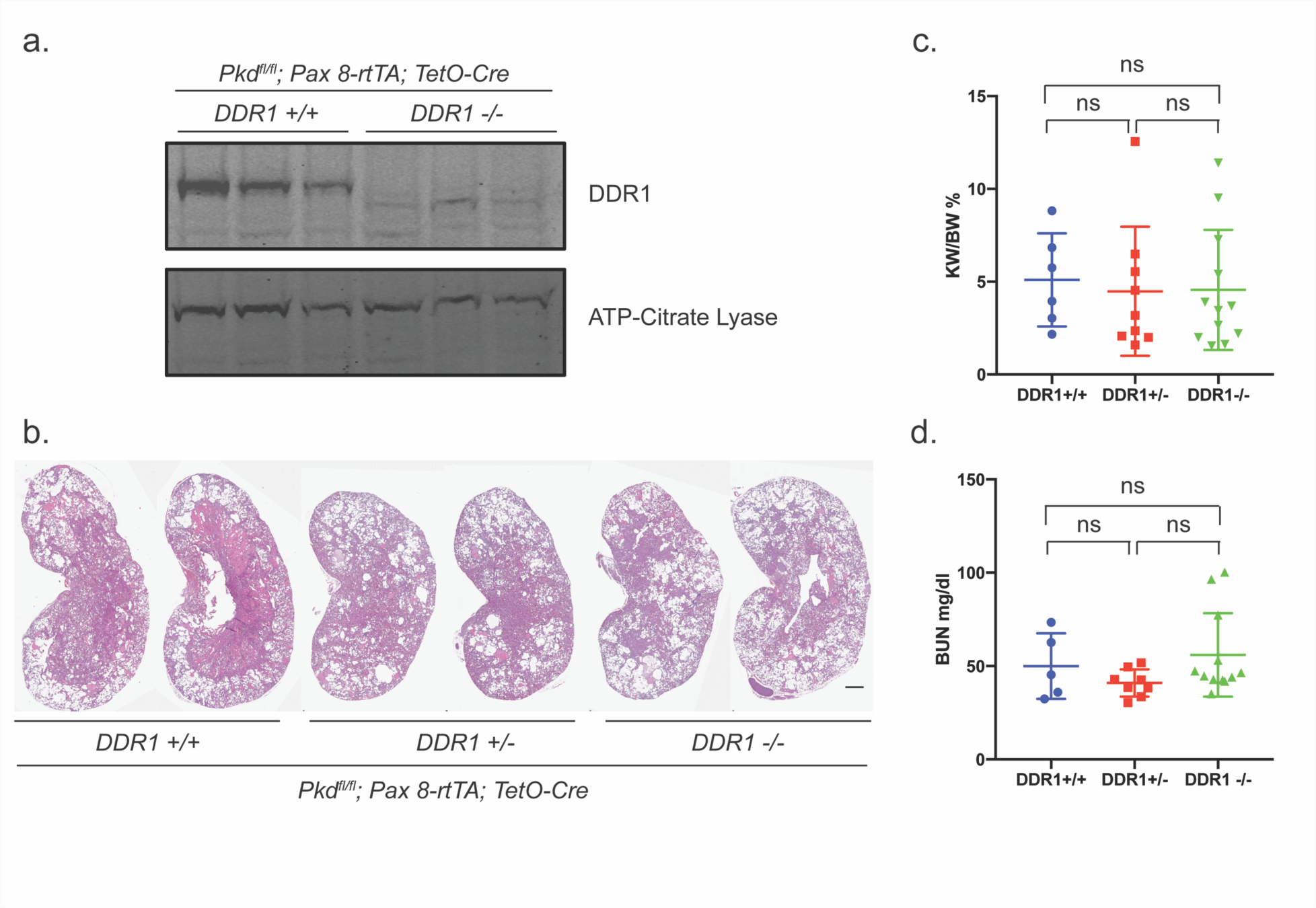
Genetic deletion of DDR1 does not slow cyst growth of preserve renal function in the inducible late onset *Pax8rtTA; TetO-Cre; Pkd1*^*fl/fl*^ mouse model. **(a)** Lysates from kidneys from *DDR1*^*-/-*^*; Pax8rtTA; TetO-Cre; Pkd1*^*fl/fl*^ and *DDR1*^*+/+*^*; Pax8rtTA; TetO-Cre; Pkd1*^*fl/fl*^ mice probed with anti-DDR1 antibodies and anti-ATP citrate lyase as a loading control. **(b)** Representative H&E staining of sagittal sections from the 3 groups of mice with kidneys from 2 independent mice of each genotype shown. **(c)** kidney/body weight ratio and **(d)** (BUN.

## Discussion

Tolvaptan is the first FDA approved drug that has been shown to slow cyst growth and preserve renal function in patients with ADPKD. However, the relative limited benefit and significant side effects of tolvaptan treatment highlights the critical need to utilize unbiased approaches to broadly screen ADPKD kidneys for new therapeutic targets to slow cyst growth and/or interstitial fibrosis in patients with ADPKD. We utilized a powerful kinome wide method to identify in an unbiased manner kinases more active in PKD kidneys when compared with WT control kidneys with the goal that identifying a more complete list of kinases relevant to cyst growth *in vivo* would offer the potential to find better and safer therapeutic drug targets to treat patients with ADPKD. In addition, a more complete picture of kinases activated in PKD and the signaling pathways they regulate should provide additional insights into the signaling hubs and networks that are aberrantly activated in PKD kidneys and provide clearer links between upstream and downstream signaling pathways.

DDR1 was one of many kinases identified in our screen that was more active in PKD kidneys and was localized to cyst lining epithelia. The relevance of DDR1 to cancer progression, its potential role in the pathogenesis of a variety of other kidney disease, together with the possibility that DDR1 would provide new insight into how extracellular matrix impacts cyst growth suggested to us that DDR1 was an exciting and promising candidate to study. However, the inability of genetically deleting DDR1 to slow cyst growth and preserve renal function in both an “early rapid” and “late slow” mouse model of PKD conclusively demonstrates that DDR1 does not play a role in PKD pathogenesis and thus is not a viable drug target.

In spite of our negative results, we think publication of our findings will be of interest and value to the PKD and nephrology community. First, these studies will prevent other researchers from potentially spending money and time assessing the relevance of DDR1 to PKD pathogenesis. Second, our studies provide an important proof of principle whereby new technologies can be used to rapidly screen for potential new drug targets, which then can be rapidly assessed *in vivo* for their relevance to disease using CRISPR/Cas9, an approach we are currently undertaking in the lab with success. Third, our findings reinforce the importance of a dasatinib-sensitive kinase(s) in PKD pathogenesis. Identifying the dasatinib specific kinase(s) that contributes to pathogenesis coupled with a drug that more specifically targets this kinase(s), could potentially lead to a safe and efficacious drug to treat patients.

## Acknowledgements

Portions of this work was supported by grants from the PKD foundation to IS and EYS.

